# Genome-wide analysis of experimentally evolved *Candida auris* reveals multiple novel mechanisms of multidrug-resistance

**DOI:** 10.1101/2020.09.28.317891

**Authors:** Hans Carolus, Siebe Pierson, José F. Muñoz, Ana Subotić, Rita B. Cruz, Christina A. Cuomo, Patrick Van Dijck

## Abstract

*Candida auris* is globally recognized as an opportunistic fungal pathogen of high concern, due to its extensive multidrug-resistance (MDR). Still, molecular mechanisms of MDR are largely unexplored. This is the first account of genome wide evolution of MDR in *C. auris* obtained through serial *in vitro* exposure to azoles, polyenes and echinocandins. We show the stepwise accumulation of multiple novel mutations in genes known and unknown in antifungal drug resistance, albeit almost all new for *C. auris*. Echinocandin resistance evolved through a codon deletion in *FKS1* accompanied by a substitution in *FKS1* hot spot 3. Mutations in *ERG3* and *CIS2* further increased the echinocandin MIC. Decreased azole susceptibility was acquired through a gain of function mutation in transcription factor *TAC1b* yielding overexpression of the drug efflux pump Cdr1; a segmental duplication of chromosome 1 containing *ERG11*; and a whole chromosome 5 duplication, which contains *TAC1b*. The latter was associated with increased expression of *ERG11, TAC1b* and *CDR2*, but not *CDR1*. The simultaneous emergence of nonsense mutations in *ERG3* and *ERG11*, presumably leading to the abrogation of ergosterol synthesis, was shown to decrease amphotericin B susceptibility, accompanied with fluconazole cross resistance. A mutation in *MEC3*, a gene mainly known for its role in DNA damage homeostasis, further increased the polyene MIC. Overall, this study shows the alarming potential and diversity for MDR development in *C. auris*, even in a clade until now not associated with MDR (clade II), hereby stressing its clinical importance and the urge for future research.

**Importance:** *C. auris* is a recently discovered human fungal pathogens and has shown an alarming potential for multi- and pan-resistance towards all classes of antifungals most commonly used in the clinic. Currently, *C. auris* has been globally recognized as a nosocomial pathogen of high concern due to this evolutionary potential. So far, this is the first study in which the stepwise progression of MDR in *C. auris* is monitored *in vitro*. Multiple novel mutations in known ‘resistance genes’ and genes previously not or vaguely associated with drug resistance reveal rapid MDR evolution in a *C. auris* clade II isolate. Additionally, this study shows that *in vitro* experimental evolution can be a powerful tool to discover new drug resistance mechanisms, although it has its limitations.

## Introduction

Over the course of a decade since its discovery (1), *Candida auris* has emerged in at least 39 countries along every inhabited continent (2), occasionally causing healthcare-associated outbreaks of lethal candidiasis (3). *C. auris* is substantially different from any other *Candida* species studied so far, as it behaves like a true multidrug-resistant (MDR) nosocomial pathogen (cfr. methicillin resistant *Staphylococcus aureus*, MRSA) (3). This was illustrated by the US Center for Disease Control and Prevention (CDC) in 2019, as they listed *C. auris* as the first fungus among urgent antimicrobial resistance threats (4). *C. auris* can become resistant to each drug and each combination of drugs from the three major antifungal drug classes: the azoles (e.g. fluconazole), echinocandins (e.g. caspofungin) and polyenes (e.g. amphotericin B). Various clinical isolate screening reports indicate fluconazole resistance in over 80% (5–9) and amphotericin B resistance in up to 30% of the isolates tested (6, 7). Echinocandin resistance is less common, found in 2-10% in some screenings (6–8, 10), but it is alarmingly on the rise (11, 12). Overall, about 90% of the *C. auris* isolates are estimated to have acquired resistance to at least one drug, while 30-41% are resistant to two drugs, and roughly 4% are pan-resistant (resistance to the three major antifungal drug classes) (4, 7). These numbers show an unprecedented potential to acquire MDR, unlike any other pathogenic *Candida* species (3, 12, 13). The molecular mechanisms of antifungal drug resistance, especially for amphotericin B resistance and MDR, are still poorly understood in *C. auris*. Hundreds of resistant *C. auris* strains have been sequenced and their decreased drug susceptibility for azoles and echinocandins has been associated with a handful of mutations in genes known to be involved in drug resistance. Still, the high levels of resistance and extensive MDR in some strains cannot be explained through the limited number of resistance-conferring mutations described so far (3, 7).

Azole resistance has been linked to three single nucleotide polymorphisms (SNPs) (7–9, 14) and an increased copy number (9, 15) of *ERG11*, the gene encoding the fluconazole target lanosterol 14-α-demethylase. Additionally, the ATP Binding Cassette (ABC) transporter Cdr1, was proven to act as an efflux pump of azoles in *C. auris* (16–18) and a recent study suggests that gain of function (GOF) mutations in *TAC1b* can underly this mode of action (16). Echinocandin resistance in *C. auris* was previously only linked to SNPs substituting amino acid S639 (9, 12, 19) and F635 (20) of Fks1, which is the echinocandin target: β(1,3) D-glucan synthase. The polyene amphotericin B works by sequestering ergosterol, rather than inhibiting a specific enzyme and therefore, amphotericin B resistance is among the least understood drug resistance mechanisms in *C. auris* and *Candida* sp. in general (12). So far, only an increased expression of genes involved in ergosterol biosynthesis (i.e. *ERG1, ERG2, ERG6* and *ERG16*) (15), SNPs in the gene encoding the transcription factor Flo8 and an unnamed membrane transporter encoding gene (21), had been linked to amphotericin B resistance in *C. auris* (12, 19). Overall, few studies have actually been able to validate the proposed drug resistance mechanisms in *C. auris* (16, 18, 22, 23), presumably because of the lack of an optimized gene-editing system for *C. auris*.

Here, we circumvent challenging and laborious gene-editing in *C. auris* (16, 22, 24) through a strategy of serial transfer based experimental evolution with the ability to trace back the emergence - and validate the causality - of single mutations or copy number changes throughout the evolutionary process. By designing allele specific PCR primers, the presence or absence of specific mutations could easily be screened for by PCR on multiple single clones in the daily evolving populations. Doing so, we validated 10 non-synonymous mutations in eight genes, evolved in 5 separate evolutionary lineages. When drug resistance became apparent, single clones from each lineage were whole genome sequenced and the SNPs, insertions/deletions (Indels) and copy number variants (CNVs) discovered were validated.

In this study we investigated MDR evolution in a clade II *C. auris* strain, which is understudied comparted to other clades (25) and has been suggested to be less prone to drug resistance development (25, 26). Previously, five different clades (clade I-V, i.e. the South Asian, East Asian, African, South American and Iranian clade resp.) of *C. auris* have been identified, each separated by thousands of SNPs (7, 27), and often associated with clade-specific virulence and/or drug resistance tendencies (9, 25, 26). As such, this study shows that clade II *C. auris* can also rapidly acquire MDR *in vitro* and its mechanisms of resistance provide fundamental knowledge on how resistance can be acquired by *C. auris*. Finally, our study presents both the power and challenges of using *in vitro* experimental evolution to discover the molecular mechanisms of (multi)drug-resistance.

## Results

### *C. auris* clade II can acquire multidrug-resistance rapidly *in vitro*

*C. auris* strain B11220, the original type strain described by Satoh *et al*. (1) in Japan 2009, was the only strain used for this study. A single colony isolate from strain B11220 (further referred to as wild type, wt), was subjected to an *in vitro* experimental micro-evolution assay as depicted and described in **figure 1A** and the methods section respectively. The wild type strain proved to be pan-susceptible (determined by minimum inhibitory concentration or MIC_50_, see method section) to the three major antifungal drug classes: fluconazole MIC_50_: 1μg/ml, caspofungin MIC_50_: 0.125μg/ml and amphotericin B MIC_50_: 0.5μg/ml. Based on these MIC_50_ values, the wild type strain was exposed (in triplicate) to three concentrations of each drug: 2xMIC_50_, 1xMIC_50_ and 0.5xMIC_50_ or no drug, representing three selective pressures and a control, respectively. Serial transfer with conditional drug treatment (**figure 1A**) was maintained for 30 days or until drug resistance became evident from regular MIC testing. An overview of the ancestry of the evolved strains is depicted in **figure 1B**.

**Figure 1.**
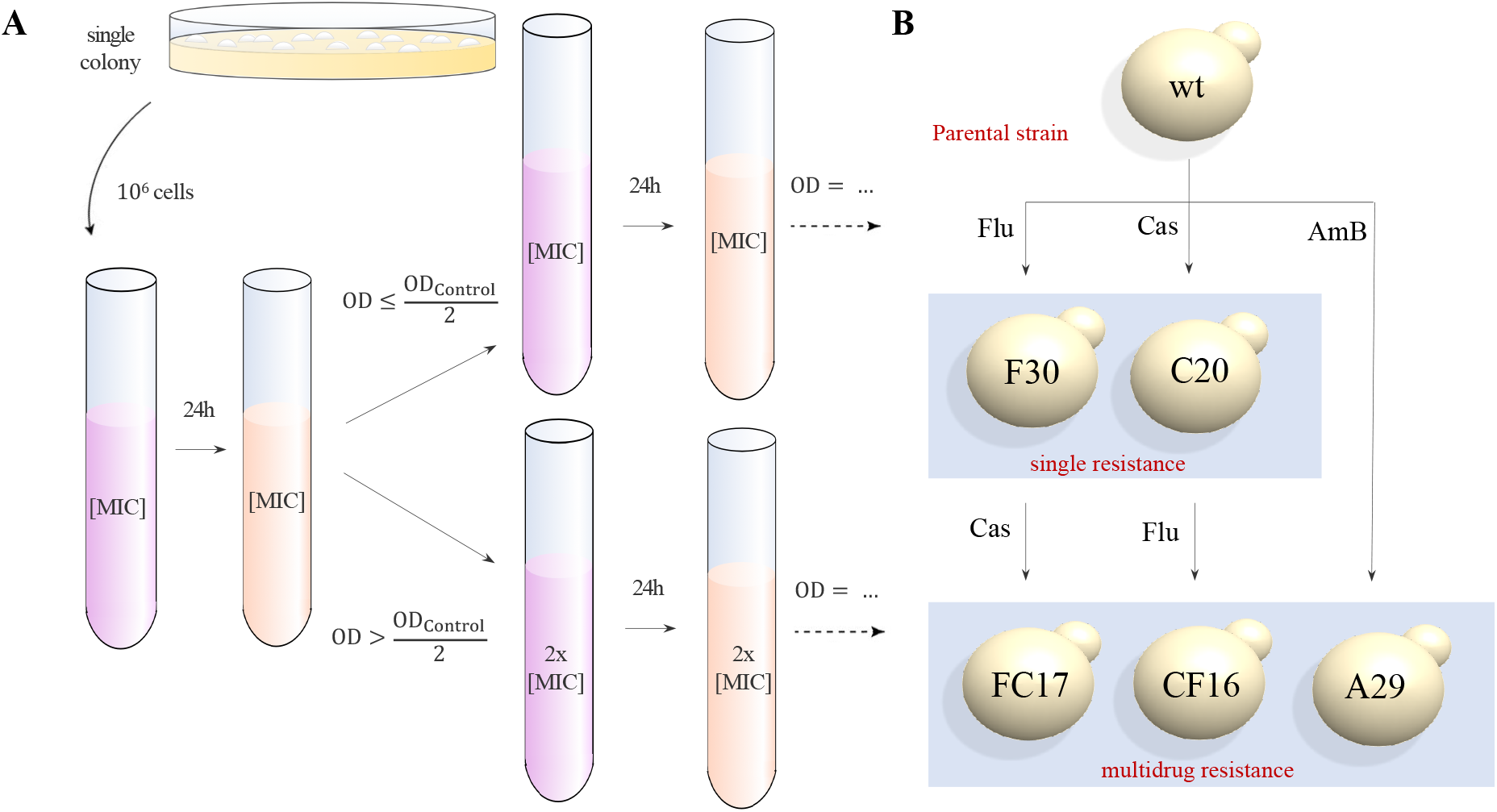
Schematic overview of the *in vitro* experimental evolution. **A)** The evolution assay: A single colony is cultured in RPMI-MOPS medium (2% glucose) for 24h at 37°C after which a standardized inoculum (10^6^ cells) is resuspended in medium containing no drug (control), the drug at a concentration of 2xMIC_50_, 1xMIC_50_ and 0.5xMIC_50_ (shown here) of the particular starting strain. Daily, the culture is re-diluted (1/10) in fresh RPMI-MOPS medium (2% glucose) with a concentration of drug based on the OD_600_ of the control culture. All strains were evolved in triplicate. Daily aliquots of evolving populations were stored in RPMI-MOPS medium containing 25% glycerol at −80°C for later examination. **B)** Ancestry of the five evolved strains that were sequenced. WGS was performed on a single colony. The name of each strain represents the experimental treatment (letter) and day of isolation (number), respectively.

Five strains were evolved and sequenced: F30, C20, A29, FC17 and CF16: strain B11220 (wild type) was exposed to **<u>F</u>**luconazole (**<u>F</u>**-lineage), **<u>C</u>**aspofungin (**<u>C</u>**-lineage) and **<u>A</u>**mphotericin B (**<u>A</u>**-lineage), after which the single resistant strains obtained were exposed to a second drug to acquire multidrug-resistance, yielding the **<u>FC</u>**- and **<u>CF</u>**-lineage for the **<u>F</u>** (**<u>F</u>**luconazole resistant) strain that was given **<u>C</u>**aspofungin and the **<u>C</u>** (**<u>C</u>**aspofungin resistant) strain that was given **<u>F</u>**luconazole respectively. The name of each strain represents the experimental lineage (letter which refers to the treatment/resistance), and day of isolation (number), respectively. **Figure 2** shows the MIC_50_ values for each drug and each end-point strain (F30, C20, A29, FC17 and CF16) evolved. The length of the evolution experiment ranged from 16 (CF-lineage) to 30 days (F-lineage), although later it was shown that resistant clones emerged quite early (e.g. after three days in C-lineage, see **figure 2**).

**Figure 2.**
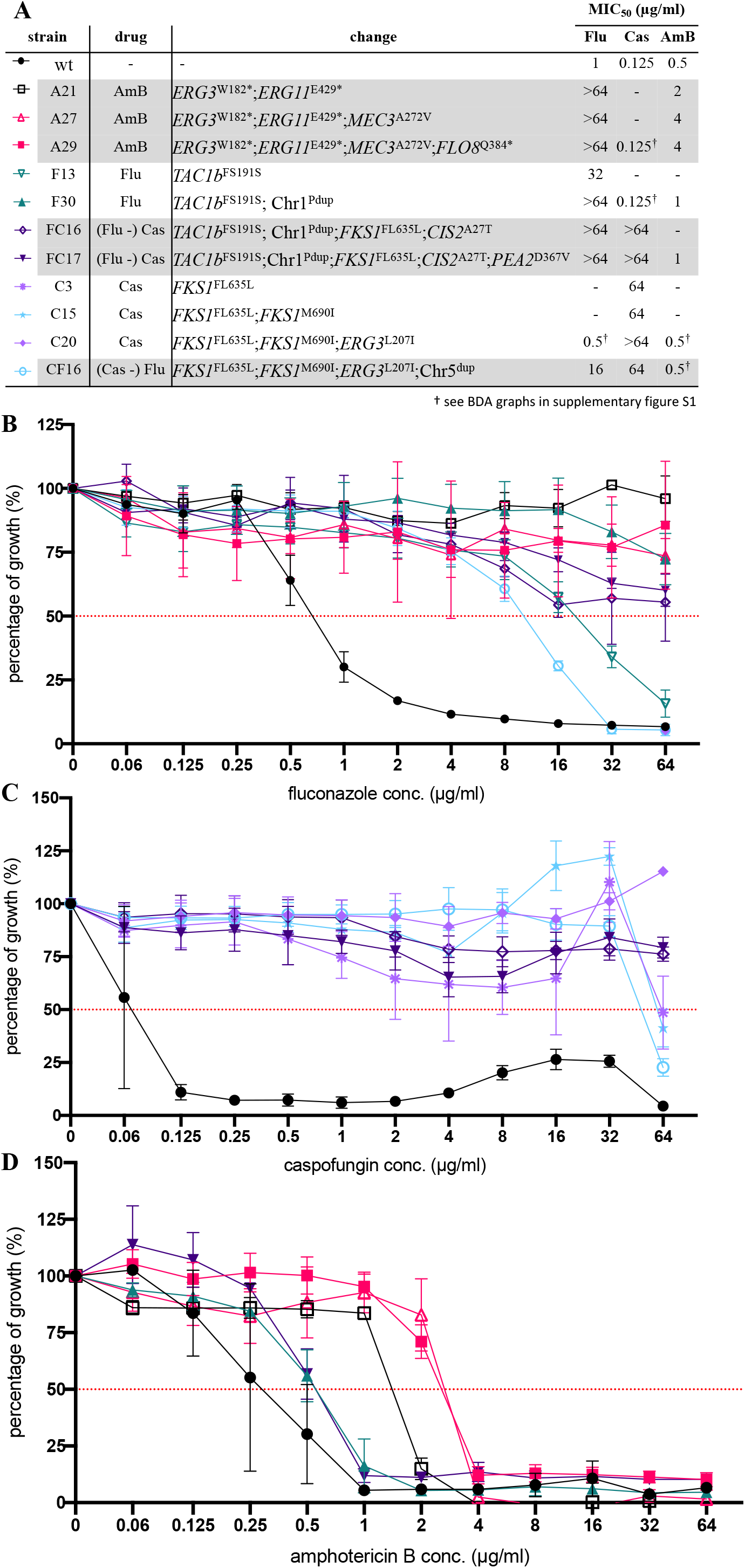
Resistance profiles of evolved strains. **A)** Summary of MIC_50_ values and associated mutations/CNVs for each end-point strain and divergent intermediate strain. **B-D)** Growth profiles of evolved strains relative to the wild type strain (wt) in a BDA of fluconazole **(B)**, caspofungin **(C)** and amphotericin B **(D)** respectively. Percentage of growth was calculated from growth without drug and based on OD_600_ measurements after 48h of incubation. Each data point and its standard deviation is calculated from 3 biological repeats, each represented by the mean of 2 technical repeats. Pdup: partial duplication, dup: duplication

### Allele-specific PCR is an effective method for tracing back the emergence of mutations during evolution in serial isolates

After micro-evolution, whole genome sequencing of the wild type strains and strain F30, C20, A29, FC17 and CF16 showed the acquisition of 10 non-synonymous mutations (listed in **table 1**) and two different aneuploidies (shown in **figure 6**, validation shown in supplementary **figure S3**). All mutations identified were novel to *C. auris* based on literature review and comparison of sequences with a set of 304 globally distributed *C. auris* isolate sequences representing Clade I, II, III and IV (9). The impact of individual mutations and CNVs will be discussed in the following paragraphs.

**Table 1.** All non-synonymous mutations identified in the end-point evolved strains. Nucleotide and amino acid changes, as compared to the reference parent strain genome (wild type) are displayed. Genes were identified based on orthologues annotated in the *C. auris* B8441 genome sequenced as provided at http://www.candidagenome.org/

| strain | change | type of change | Gene ID (B11220) | Gene ID (B8441) | Ortholog |
| --- | --- | --- | --- | --- | --- |
| **A29** | tgG/tgA\|W182* | nonsense | CJI96_002270 | B9J08_003737 | *ERG3* |
|  | Gag/Tag\|E429* | nonsense | CJI96_001197 | B9J08_001448 | *ERG11* |
|  | Cag/Tag\|Q384* | nonsense | CJI96_001121 | B9J08_000401 | *FLO8* |
|  | gCg/gTg\|A272V | missense | CJI96_001637 | B9J08_003102 | *MEC3* |
| **F30** | ttcagt/agt\|FS191S | codon deletion | CJI96_004335 | B9J08_004820 | *TAC1b* |
| **FC17** | ttcagt/agt\|FS191S | codon deletion | CJI96_004335 | B9J08_004820 | *TAC1b* |
|  | ttcttg/ttg\|FL635L | codon deletion | CJI96_001351 | B9J08_000964 | *FKS1* |
|  | gAt/gTt\|D367V | missense | CJI96_001286 | B9J08_001356 | *PEA2* |
|  | Gca/Aca\|A27T | missense | CJI96_001769 | B9J08_003232 | *CIS2* |
| **C20** | atG/atA\|M690I | missense | CJI96_001351 | B9J08_000964 | *FKS1* |
|  | ttcttg/ttg\|FL635L | codon deletion | CJI96_001351 | B9J08_000964 | *FKS1* |
|  | Cta/Ata\|L207I | missense | CJI96_002270 | B9J08_003737 | *ERG3* |
| **CF16** | atG/atA\|M690I | missense | CJI96_001351 | B9J08_000964 | *FKS1* |
|  | ttcttg/ttg\|FL635L | codon deletion | CJI96_001351 | B9J08_000964 | *FKS1* |
|  | Cta/Ata\|L207I | missense | CJI96_002270 | B9J08_003737 | *ERG3* |

To validate the causality of single mutations in strains that harbored more than one mutation, we applied a screening strategy of allele-specific PCR (AS-PCR). AS-PCR primers were designed as described by Liu *et al*. (28), implementing a specific mismatch at the third position of the 3’ end of the allele-specific primer to increase specificity. An overview of the universal, and allele specific primers used to perform AS-PCR, is given in supplementary **table S1**. The specificity and sensitivity of all AS-PCR primers was assessed by performing temperature gradient PCRs on serial dilution reference DNA template (for one example see supplementary **figure S2**). Populations were re-cultured from the −80°C collection of daily stored aliquots (populations) and AS-PCR was performed on gDNA extracted from a maximum of 30 single clones of each (daily) population. After confirmation of the emergence of a mutation of interest, alleles were verified by sequencing a +-1000 bp region spanning the allele of interest. Primers used for PCR and sequencing are given in supplementary **table S1**. Next, the influence of this single mutation on the drug susceptibility was determined by performing a broth dilution assay (BDA, see Antifungal susceptibility testing in method section). **Figure 2** shows the impact of each individual mutation on the MIC for the drug of interest for each lineage evolved, except for the A-lineage, in which the mutations in *ERG3* and *ERG11* (**table 1**) was present or absent simultaneously in the 30 clones that were checked per population.

### Novel mutations in *FKS1* and *ERG3* yield extensive echinocandin resistance with minor growth discrepancies

Caspofungin resistance was evolved twice in this study, once as monoresistance in the C-lineage, and once as multidrug-resistance in the FC-lineage, derived from the fluconazole resistant strain F30 (**figure 1B**). The susceptibility to caspofungin decreased drastically in both strain C20 and FC17 (MIC_50_ >64μg/ml) (**figure 2**). Whole genome sequencing revealed three mutations in the C20 strain: a missense mutation (atG/atA|M690I) and codon deletion (ttcttg/ttg|FL635L) in *FKS1* (B9J08_000964; **table 1**), the gene encoding the catalytic subunit of the echinocandin drug target β(1,3) D-glucan synthase, and one missense mutation (Cta/Ata|L207I) in *ERG3*, encoding sterol Δ5,6-desaturase (B9J08_003737; **table 1**). The exact same codon deletion (ttcttg/ttg|FL635L) in *FKS1* emerged independently during caspofungin resistance evolution in the FC-lineage (**table 1**). Two additional mutations emerged during the evolution of strain FC17, namely a missense mutation (gAt/gTt|D367V) in the *PEA2* gene, encoding a subunit of the polarisome (B9J08_001356; **table 1**), and a missense mutation (Gca/Aca|A27T) in the *CIS2* gene, encoding a γ-glutamylcysteine synthetase (B9J08_003232; **table 1**). Tracing back the emergence of these mutations shows that the *FKS1* mutation FL635L increased the MIC_50_ of the wild type strain 500-fold: from 0.125μg/ml to 64μg/ml (**figure 2**). It is however the mutations in *CIS2* (emerged in FC16) and *ERG3* (present in C20) that further increased the caspofungin MIC_50_ to >64μg/ml (**figure 2**).

Acquired echinocandin resistance in fungi has been associated with several specific mutations in three defined ‘hot spot’ regions (HS) in the *FKS1* gene (29). **Figure 3** shows an amino acid sequence alignment of the *FKS1* gene HS1, HS2 and HS3 regions, constructed to compare the mutations found in this study to those known to confer echinocandin resistance in *C. auris* and other fungi as described in literature. This literature review shows that the codon deletion at position F635 as found in this study; also has been reported to confer decreased echinocandin susceptibility in *C. glabrata* (29). The same amino acid was substituted (not deleted as in C-lineage here) in echinocandin resistant *C. auris* strains reported recently (20). The *FKS1* mutation M690I is located in hot spot region 3 without comparable mutations in pathogenic fungi (**figure 3**), and seems to have no direct impact on the drug susceptibility to caspofungin in the C-lineage (**figure 2**).

**Figure 3.**
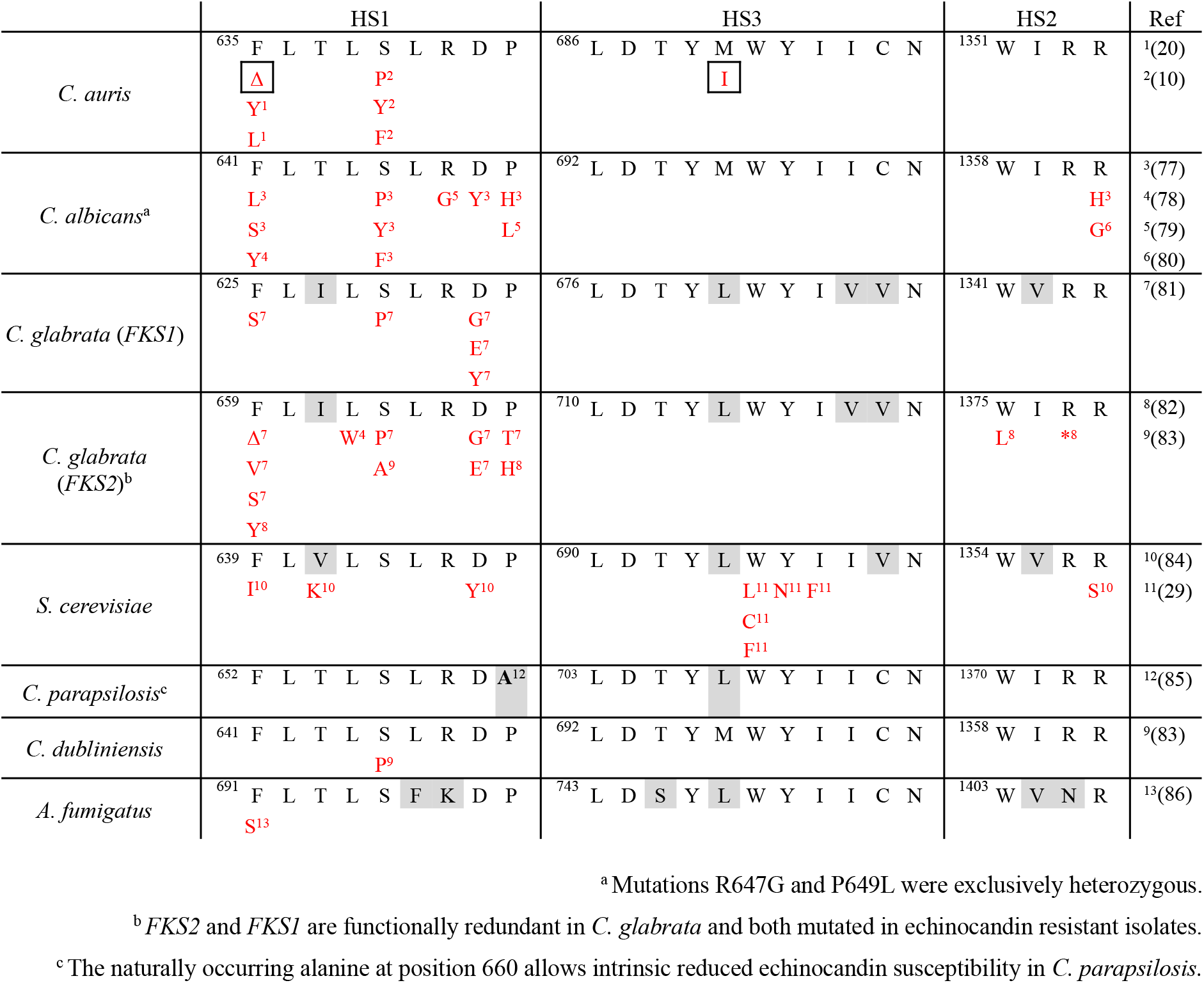
Hotspot (HS) region mutations of the *FKS* genes that confer echinocandin resistance. Amino acid sequence of hotspot 1, 2 and 3 (HS1-3 resp.) of *C. auris* and other fungi are aligned along with all mutations found to decrease echinocandin susceptibility as described in literature (references are given between brackets). Species specific polymorphisms of HS are indicated in grey, the mutations found to confer echinocandin resistance in this study are indicated by a grid. Δ: deletion, *: nonsense mutation

**Figure 4** shows the growth curves of all end-point evolved strains, plotted based on growth in RPMI-MOPS medium supplemented with 0,2% or 2% glucose (**figure 4A and 4B** respectively). From these growth plots, it is clear that caspofungin resistant strains C20 and FC17 hardly had any growth discrepancies compared to the parent wild type strain under physiological conditions, showcasing the lack of fitness trade-offs associated with the acquisition of echinocandin resistance described above.

**Figure 4.**
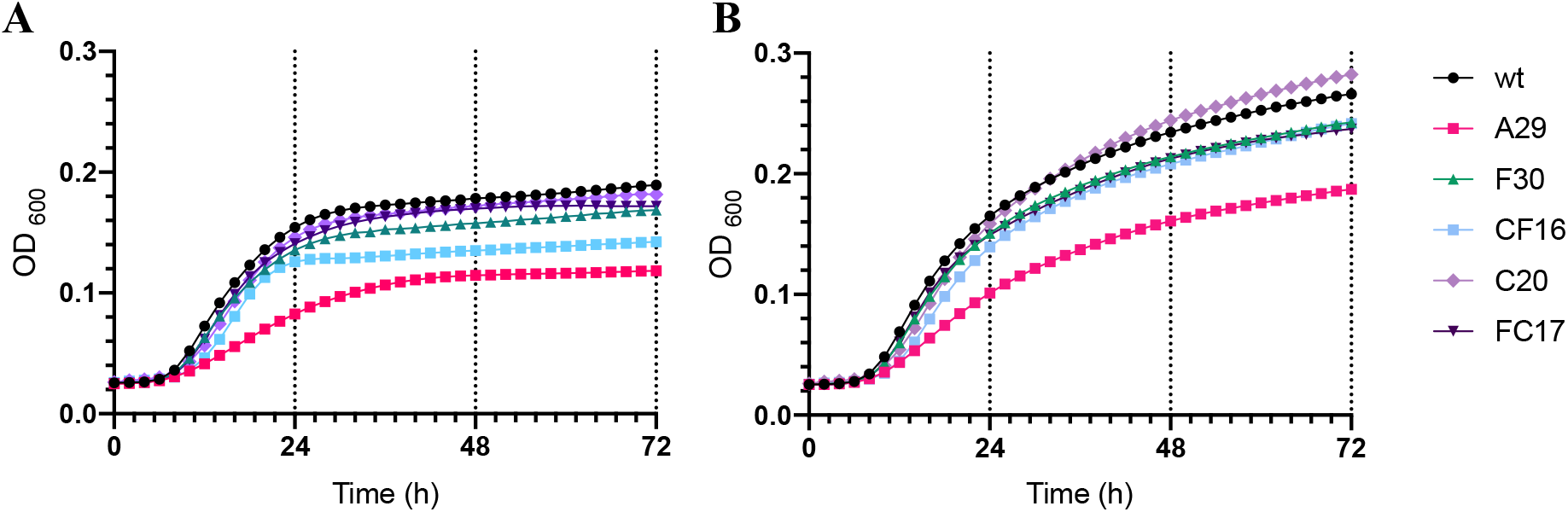
Growth curves of end-point evolved strains. Growth curves were plotted based on culture density (spectrophotometric quantification of OD_600_ see method section) over 72h of incubation in RPMI-MOPS medium containing **A)** 0.2% glucose and **B)** 2% glucose at 37°C. Data points are average values of three biological repeats represented each by the average of two technical repeats.

### Mutations of *ERG3, ERG11, FLO8* and *MEC3* results in amphotericin B and fluconazole resistance but impede growth

During micro-evolution, the MIC of amphotericin B increased 8-fold in the A-lineage: from IC_50_: 0.5μg/ml (wt) to MIC_50_: 4μg/ml in strain A29 (**figure 2**). Simultaneously, cross-resistance to fluconazole emerged, with an MIC increase from 1μg/ml to over 64μg/ml (**figure 2**). Two nonsense mutations in genes involved in the ergosterol biosynthesis pathway were found (**table 1**), namely the tgG/tgA|W182* mutation in the *ERG3* gene and the Gag/Tag|E429* mutation in the *ERG11* gene, encoding lanosterol 14-alpha-demethylase (B9J08_001448; **table 1**). The *ERG11* mutation of strain A29 lies within a region of *ERG11* that corresponds to a frequently mutated (‘hotspot’) region of *ERG11* in azole resistant *C. albicans* (30, 31). It is however distinct from the three SNPs of *ERG11* (namely Y132F, K143R and F126L) that have been linked to drug (azole) resistance in *C. auris* so far (7–9, 14), and are situated in another ‘hotspot’ region of *ERG11* (30, 31).

Additionally, a nonsense mutation (Cag/Tag|Q384*) was found in the transcription factor *FLO8* gene (B9J08_000401, **table 1**), and a missense mutation (gCg/gTg|A272V) emerged in the *MEC3* gene which encodes a subunit part of the Rad17p-Mec3p-Ddc1p sliding clamp (B9J08_003102; **table 1**). Remarkably, the mutation in *MEC3* increased the amphotericin B resistance two-fold (from MIC_50_:2μg/ml in strain L21 to MIC_50_:4μg/ml in strain L27, see **figure 2**). The mutation in *FLO8* did not seem to alter the drug susceptibility for fluconazole or amphotericin B. Additionally, strain A29 was found to significantly overexpress *TAC1b* and *ERG11*, as shown by reverse transcriptase quantitative PCR (RT-qPCR), shown in **figure 5**. Observations in the lab and characterization of the growth curve showed that strain A29 was most impeded in growth, compared to the wild type strain and other evolved strains, as shown in **figure 4**.

**Figure 5.**
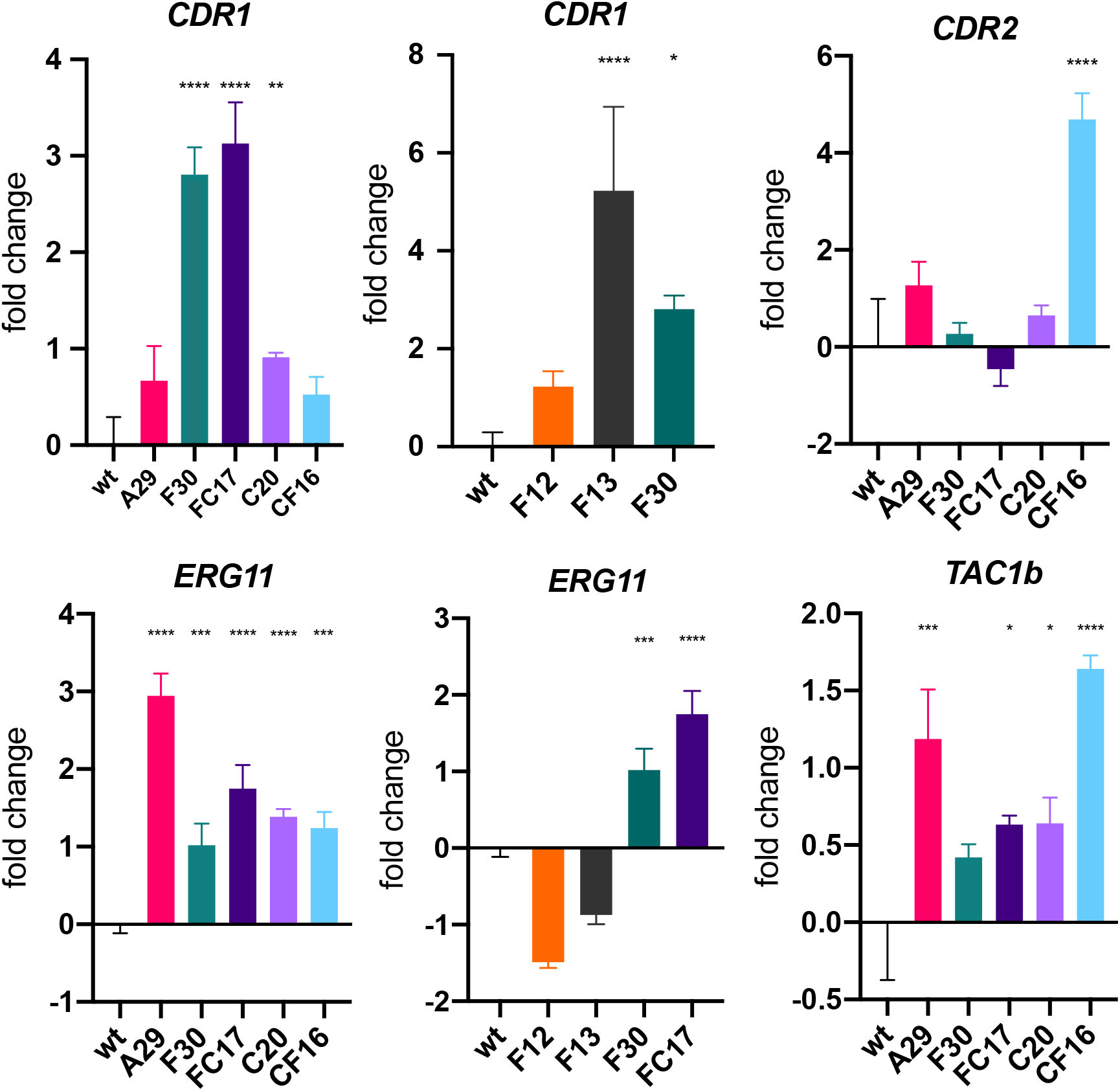
Relative expression of various genes of interest among evolved strains. Fold change of expression levels for *CDR1, CDR2, ERG11* and *TAC1b* for the wild type (wt), end-point evolved strains (A29, F30, FC17, C20, CF16) and intermediate strains F12 and F13 (for *CDR1* and *ERG11*). Bars represent log2-transformed means with standard deviation accounting for data obtained from 3 biological repeats, each represented by the mean of 2 technical repeats. Asterisks indicate significant overexpression: \**P*<0.05; \*\**P*<0.01;\*\*\**P*<0.001;\*\*\*\**P*<0.0001

### A *TAC1b* mutation and upregulated *CDR1* expression decrease fluconazole susceptibility

In strain F13, a codon deletion (ttc/|F15) in the *TAC1b* gene was identified (B9J08_004820; **table 1**) that corresponded to a 32-fold increase in the MIC_50_ of fluconazole (**figure 2**). Tac1b is an activating transcription factor, positively regulating the expression of the ATP Binding Cassette (ABC) transporter Cdr1, known to be involved in azole efflux and azole resistance in *C. auris* (16, 22). Although the mutation in strain F13 is novel, it is located in a region of *TAC1b* that is known to harbor gain-of-function mutations in fluconazole resistant clinical isolates of *C. auris*, as shown by Rybak *et al*. (16). Nevertheless, it is the first codon deletion (cfr. SNPs) in this region suggested to confer a gain of function of *C. auris TAC1b*. Gene expression analysis of strain F12 (no *TAC1b* mutation) and strain F13 (*TAC1b* mutation obtained) confirms that this mutation increased the expression of *CDR1* (B9J08_000164) significantly, as shown in **figure 5**. The overexpression of *CDR1* is maintained in strain F30 and in the multidrug-resistant strain FC17, as shown in **figure 5**.

### Two aneuploidies independently emerged during fluconazole resistance evolution

Read coverage of whole genome sequencing was used to analyze copy number variation (CNV), by calculating normalized depth read coverage per 5kb window (see methods section). A visual representation of this normalized coverage for each chromosome in all end-point sequenced strains is displayed in **figure 6**. This reveals a segmental and whole chromosome duplication emerged in the F- and CF-lineage respectively, both seemingly involved in fluconazole resistance evolution.

**Figure 6.**
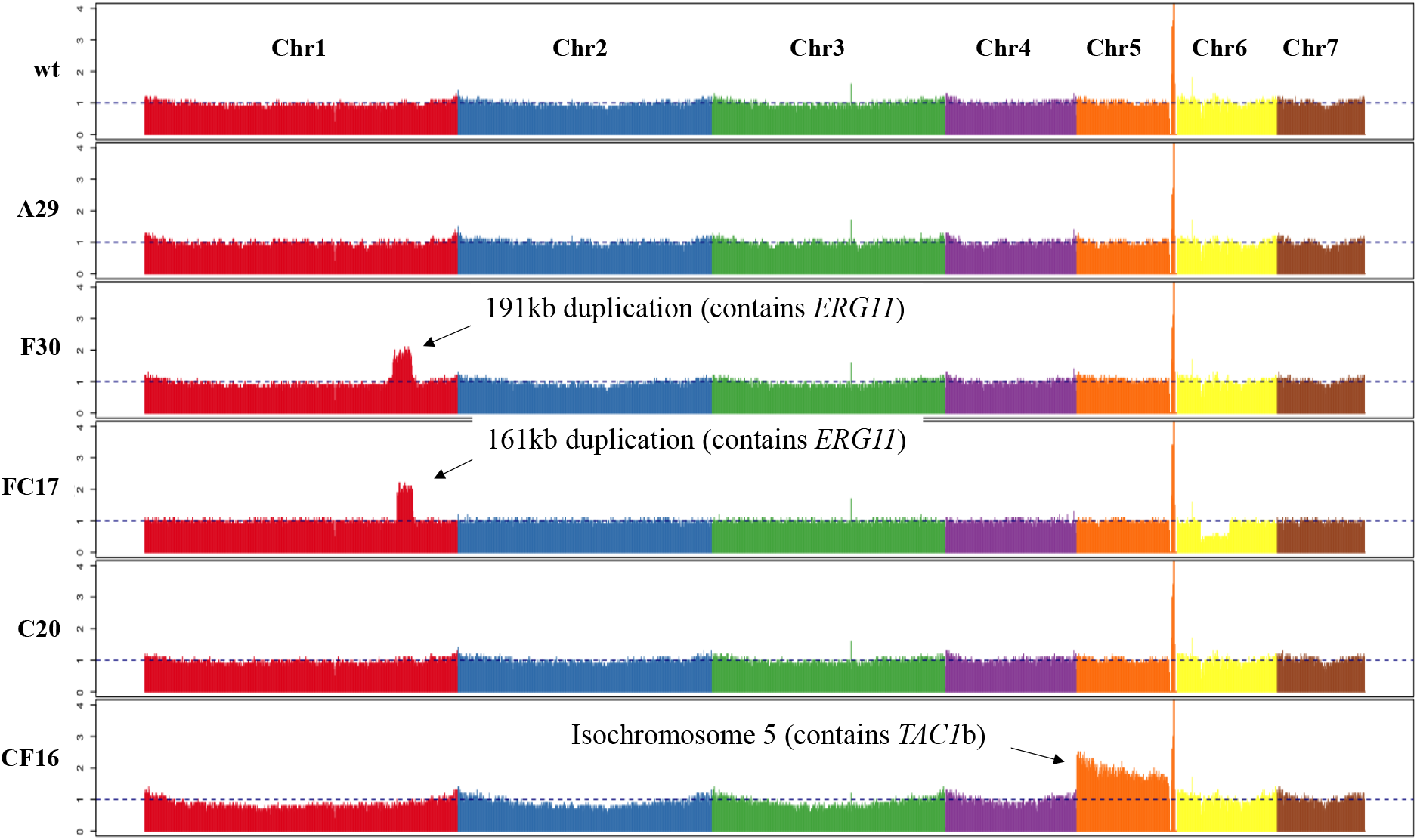
Coverage plot of whole genome sequencing of end-point evolved strains. The coverage displayed is calculated by normalizing the average coverage depth per 5kb window. Each color represents 1 chromosome (f.l.t.r.: chromosome 1 to 7). Indicated are the significant duplication in chromosome 1 (Chr1) in strain F30 and FC17 and chromosome 5 (Chr5) in strain CF16. Anomality’s in Chr5 (spike in all strains) and Chr6 (decreased coverage) are caused by ribosomal DNA (rDNA) and unambiguous mapping, and can thus be ignored.

The 191 kb segmental duplication of Chr1 in the F30 strain contained 75 protein encoding genes (based on the B11220 reference genome annotation; CP043531-CP043537), including *ERG11*. During further evolution to caspofungin in the FC-lineage (see **figure 1B**), this segmental duplication was maintained but decreased in size to 161kb containing 67 protein encoding genes (still including *ERG11*). The segmental Chr1 duplication resulted in an over two-fold decrease in fluconazole susceptibility, increasing the MIC_50_ of 32μg/ml in strain F13 to MIC_50_ >64μg/ml in strain F30, as shown in **figure 2**. Expression analysis showed that the duplication led to increased expression of *ERG11*, not present in strain F13 (**figure 5**). Strain F30 was also marked by a slight decrease in amphotericin B susceptibility, retained in the FC17 strain (**figure 2**). This is possibly due to *ERG11* overexpression.

The whole chromosome 5 (Chr5) duplication in the CF16 strain contained a region of 933kb encompassing 405 protein encoding genes, including and *TAC1b*. This aneuploidy marks the difference between strain C20 and strain CF16 and is therefore suggested to confer the 32-fold decrease in fluconazole susceptibility between those strains (**figure 2**). Expression analysis showed that the duplication of Chr5 correlates with a significant overexpression of *TAC1b* and *CDR2* (B9J08_002451), but not *CDR1* in strain CF16 (**figure 5**).

## Discussion

### First, this study shows the evolution of multiple mechanisms known to be involved in drug resistance in fungi, albeit new to *C. auris*

In the largest screening of *C. auris* clade II strains, 62.3% of a total of 61 isolates proved to be fluconazole resistant although only 3 isolates harbored a known azole-conferring mutation in *ERG11* (K143R) (32). This indicates that other mechanisms of azole resistance play a role in *C. auris* clade II (32). Here, we describe at least four molecular mechanisms, none of which include the most common mutations in *ERG11*, by which fluconazole susceptibility can decrease in a clade II *C. auris* strain *in vitro*. Previous reports show that many GOF mutations in *TAC1* or homologues of this transcription factor can confer azole resistance in *Candida* species (33–35), including *C. auris* (16), through an overexpression of the drug efflux pump Cdr1. Most GOF mutations are found in the region encoding the putative transcriptional activation domain of *TAC1*, situated in the C-terminal portion of the protein in *Candida* sp. (36). Rybak *et al*. report one mutation in this region (codon deletion at position F862) to be associated with fluconazole resistance in *C. auris* although all other resistance associated mutations lie between the DNA binding, transcription factor and activation domain of *TAC1b* (16). One such mutation (F214S), discovered in an experimentally evolved strain of *C. auris*, lies in the proximity of the codon deletion at position 191 as we discover here. Based on these reports and our findings, we therefore hypothesize that F191Δ is a new potential gain of function mutation of *C. auris TAC1b* conferring azole resistance through *CDR1* overexpression. Nevertheless, previous reports have shown that Tac1b might function in other, Cdr1-independent ways to decrease azole susceptibility in *C. auris* (16, 22). Another mechanism of reduced azole susceptibility discovered in this study involves aneuploidies. In *C. albicans*, both *TAC1* and *ERG11* are located on Chr5, and the duplication of this region by forming an isochromosome [i(5L)] has been reported to confer azole resistance *in vitro* and *in vivo* (37). Based on this and reports on azole resistance in *C. auris* due to CNVs and/or overexpression of *ERG11* (9, 15, 38), we hypothesize a similar mode of action in strain F30. Moreover, comparing drug susceptibility between strain F13 and F30, the overexpression of *ERG11* in F30 more than doubles the fluconazole MIC_50_ compared to-the already resistant-strain F13, while it decreases the susceptibly for amphotericin B (**figure 2**). Given the fact that the duplication of *C. auris* Chr5 is the only genomic alteration that distinguishes strain CF16 from strain C20, we propose that this duplication is here too responsible for azole resistance. Expression analysis shows however, that the duplication and subsequent overexpression of *TAC1b* (**figure 5**) does not correspond to an increased expression of *CDR1*, as was expected, but *TAC1b* may play a CDR1-independent role in azole resistance of *C. auris*, as suggested by Mayr *et al*. (22).

The acquisition of resistance to polyenes is among the least understood of all antifungal drugs and has been linked to mutations in the ergosterol biosynthesis pathway in *Candida* sp. including *ERG2* (39), *ERG6* (40), *ERG11* (41) and *ERG3* (42). Cross resistance to azoles and amphotericin B has often been associated to the abrogation of two *ERG*-genes simultaneously (43). One such example is the combination of the loss of function (LOF) of *ERG11* and *ERG3* in *C. tropicalis* (44, 45). Upon the abrogation of *ERG11*, due to a LOF mutation or the action of azoles, a toxic 3,6-diol derivative is produced through the action of the sterol Δ^5,6^ desaturase, encoded by *ERG3* (46). Simultaneous disruption of the function of both *ERG3* and *ERG11* can undo this detrimental effect (43). Here we show for the first time that such a mechanism of cross-resistance can establish in *C. auris*.

Target alteration is the most commonly observed and most studied mechanism of echinocandin resistance in *Candida* species (47). Most echinocandin resistance conferring *FKS1* mutations in *C. auris* occur at position S639 (12, 19) although most recently, a SNP at position F635, the same codon deleted in strain C3 in this study, was linked to resistance in the clinic (20), see **figure 3**. In general, mutations in echinocandin resistant *Candida* species lie within two small, strictly defined ‘hot spot regions’ of *FKS1* (47). However, the codon substitution at position 690, emerged in strain C15 occurs in the elusive ‘hot spot 3’, a third potent hot spot region discovered by site-directed mutagenesis of *S. cerevisiae* (29). This mutation occurred after the codon deletion at position 635 (in HS1) in the C-lineage but did not affect the echinocandin MIC_50_, possible indicating functional compensation of the altered Fks1 protein. A third mutation of strain C20 occurred in *ERG3*. One report shows that a mutation in *ERG3* in a clinical *C. parapsilosis* strain conferred both resistance to azoles and echinocandins (17). Here we observe a slight increase, rather than a decrease in fluconazole susceptibility upon the emergence of the *ERG3* mutation in strain C20 (**figure 2**). Most interestingly this mutation further increases the MIC_50_ for caspofungin in strain C20, compared to strain C15, which only obtained *FKS1* mutations (**figure 2**). Overall, the caspofungin resistant strains (FC17, C20) show MIC values (>64μg/mL) that exceed previously reported values in *C. auris* (8, 48–50). Like Rybak *et al*. (17), we therefore suggest that the underlying mechanisms of echinocandin resistance in *C. auris*, including the role of *ERG3*, should be further investigated.

### Four genes were mutated that were previously not or vaguely associated with drug resistance in fungi

*FLO8*, mutated in the amphotericin B resistant strain A29, encodes a transcription factor known to be essential for filamentation *C. albicans* (51). This filamentation was shown to decrease the rate of programmed cell death in *C. albicans*, when exposed to amphotericin B (52). Flo8 has multiple downstream effects, one of which is the positive regulation of *ERG11* expression shown in *S. cerevisiae* (53) and thus potentially playing a role in azole and amphotericin B resistance. In a recent study of clinical *C. auris* isolates from South-America, a non-synonymous mutation in the *FLO8* gene significantly correlated to amphotericin B resistance (21). In a follow-up study on the structure of FLO8, the authors suggest a potential role of FLO8 in *C. auris* virulence and drug resistance, arguing that the *FLO8* mutation found before (21), could be a gain of function mutation (54). In our study however, we see a nonsense mutation, abrogating Flo8 at amino acid 100, assuming to be disruptive to its function. Earlier, a LOF of *FLO8* was found to play a role in azole resistance with a *FLO8* deletion correlated to increased *TAC1, CDR1* and *CDR2* expression and azole resistance, while *FLO8* overexpression lead to decreased *CDR1* expression (55). Although these reports strengthen the suspicion of a role of Flo8 in drug resistance, we cannot validate the influence on the resistance phenotype of the nonsense mutation observed here. Further research on Flo8 in drug resistance is therefore highly desirable.

The fourth gene mutated during amphotericin B resistance evolution is an ortholog of *MEC3*, encoding a DNA damage checkpoint protein as part of the Rad17p-Mec3p-Ddc1p sliding clamp, involved primarily in DNA damage recognition and repair in *S. cerevisiae* (56). No clear reports of a function for *MEC3* in antifungal drug resistance were found, although two studies mention the upregulation of *MEC3* upon the acquisition of azole resistance in an experimentally evolved *C. glabrata* strain (35, 57). Our results show that the mutation in *MEC3* has a significant influence on susceptibility to amphotericin B, doubling the MIC_50_ (**figure 2**). The mechanism behind increased amphotericin B resistance upon acquiring a mutation in *MEC3*, remains unclear.

Strain FC17 harbored a mutation in *CIS2*, of which the *S. cerevisiae* ortholog (*ECM38*) encodes a γ-glutamyltranspeptidase, involved in glutathione degradation (58), detoxification of xenobiotics (59), and cell wall biogenesis (60). The role *CIS2* plays in the latter, regarding echinocandin resistance, remains unclear but as for the former, a study from Maras and colleagues (61) illustrated that fluconazole and micafungin resistance was accompanied by altered levels of glutathione in *C. albicans*, hypothesized to counteract oxidative stress caused by these antifungal drugs. In that study, the increased levels of glutathione were accompanied by the overexpression of γ-glutamylcysteine synthetase (61). A role for *CIS2* and glutathione catabolism, in drug resistance mediated by an altered redox metabolism remains to be elucidated.

The fourth mutation in the FC-lineage lays within a gene predicted to encode *PEA2*, a subunit of the polarisome, involved in polarized growth and morphogenesis in *S. cerevisiae* (62). This mutation has however no significant effect on the drug resistance profile and might thus be the result of random genetic drift.

### Experimental evolution can be a powerful tool to research resistance, although it has limitations

Due to recent advances in next generation sequencing technology, genome-wide studies of drug resistance have become more common (63, 64). The classic approach of sequencing drug resistant clinical isolates directly from patients (63) has many limitations, including the often unavailability of the original genotype and the difficulty to resolve mutations associated with drug resistance from those that have accumulated due to host-pathogen interactions. *In vitro* experimental evolution copes with most of these problems (63, 65), is highly repeatable and allows controlled long term monitoring of different strains and conditions. Moreover, the ability to isolate and investigate each generation separately, allows to monitor both the speed and the stepwise progression of drug resistance development. Nevertheless, *in vitro* experimental evolution has its own limitations, such as the homogeneity of the selective pressure in the absence of metabolization of the drug, tissue specific exposure and host-pathogen interactions. However, studies of antifungal drug resistance by *in vitro* evolution often resemble acquired resistance found in clinical isolates (63, 65). In regards to our results, a comparative analysis of mutations reported in literature and re-analysis of variants predicted in 304 sequenced clinical isolates of *C. auris* (9), show that most mechanisms of drug resistance proposed here are novel. One must be careful by redeeming these findings to be nonrelevant to the *in vivo* setting or clinical environment, reports on resistance mutations (providing whole genome analysis) is still scarce and the data base of 304 sequenced clinical isolates of *C. auris* (9) is still limited, with only 23% of isolates reported multidrug-resistant and only include 7 clinical isolates that belong to clade II (6 isolates pan-susceptible, 1 isolate fluconazole resistant) (9). This and other studies in bacteria (66) and fungi (13, 65, 67) show that *in vitro* experimental evolution can be a powerful tool, especially if combined with an effective approach to trace the full evolutionary history of mutation events, as we did here using allele-specific PCR screening

### *Candida auris* clade II is, like most other *C. auris* clades, exceptionally capable of MDR development

Clade II *C. auris* is one of the least studied clades (25) and this report aids to close this research gap. In general, few phenotypic differences have been characterized that differentiate the five *C. auris* clades from one another, although collected isolates vary substantially in the frequency of drug resistance between clades and mutations identified (9, 15). More research should be performed identify clade specific phenotypes or evolutionary tendencies, to anticipate on geographically defined emergence and outbreaks in the field. Overall, *C. auris* is still significantly understudied, compared to other *Candida* species such as *C. albicans* and *C. glabrata*, despite the fact that it is an urgent antimicrobial resistant threat (4). Further fundamental knowledge on how *C. auris* can thrive as “nosocomial MDR fungus” is highly needed to efficiently tackle this pathogen in the future.

## Materials and methods

### Strains and growth conditions

All experiments were performed with *C. auris* strain B11220 (CBS10913) from the Westerdijk Fungal Biodiversity Center (wi.knaw.nl/). Strains were grown on YPD agar (2% glucose) at 37°C and enriched in RPMI – MOPS liquid medium containing 2% glucose at 37°C in a shaking incubator overnight. All strains, including daily aliquots of serially transferred populations in the micro-evolution assay, were stored at −80°C in RPMI – MOPS medium containing 25% glycerol.

### Antifungal susceptibility testing

The Minimal Inhibitory Concentration (MIC) was determined by a broth dilution assay (BDA) according to Clinical and Laboratory Standards Insititute (CLSI) guidelines (68). In short, a dilution of 64μg/ml to 0.06 μg/ml of each drug was prepared in RPMI-MOPS medium in a 96-well polystyrene microtiter plate. A standardized amount of 100 to 500 cells was dissolved in a final volume of 200μL per well and plates were incubated at 37°C. Growth was measured after 48h of incubation through spectrophotometric quantification of OD_600_ in a SPECTRAmax^®^ Plus 384 microplate reader (Molecular Devices). Minimal Fungicidal Concentration (MFC) was determined by spotting the resuspended 96-well microtiter plate content on YPD agar (2% glucose) and incubation for 48h. Resistance is determined through tentative breakpoints provided by the CDC (6).

### In vitro experimental evolution assay

An overview of the design of the experimental evolution assay is given in figure 1A. At the start of the evolution experiment, 10^6^ cells are diluted in a 5ml volume of RPMI-MOPS medium (2% glucose) containing no drug (control) or a drug at a concentration of 1/2xMIC_50_, 1xMIC_50_ or 2xMIC_50_. All conditions were performed in triplicate (3 evolving populations per condition). After 24h of incubation at 37°C in a shaking incubator, growth of each population was compared to the average growth of 3 controls (no drug) by spectrophotometric quantification (OD_600_). Next, 500μL of each population was transferred to 4500μL of fresh medium with a concentration of drug equal to the previous culture when OD_600_ [evolving population] ≤ OD_600_ [average control] or double to the previous culture when OD_600_ [evolving population] > OD_600_ [average control]. The experiment was terminated after 30 days or if the MIC_50_ exceeded the resistance breakpoint value, as evaluated by intermediate MIC testing.

### Analysis of growth

Growth was assessed by spectrophotometric observation (OD_595_) over time in a Multiskan™ GO (Thermo Scientific) automated plate reader using flat bottom 96-well plates and intermitted (30 min. interval) pulsed shaking (medium strength, 5 min). Cultures were diluted in RPMI-MOPS medium containing 0.2% or 2% glucose, to a final volume of 10^6^ cells per well. Growth was measured for 72h at 37°C. Growth curves were plotted as an average values of 3 biological repeats with 3 technical repeats per biological repeat.

### DNA extraction

Genomic DNA for whole genome sequencing was extracted using the MasterPure™ Yeast DNA Purification kit (Lucigen, US) following the manufacturers protocol. For (AS-)PCR and sanger sequencing, DNA was isolated from the cells through phenol chloroform isoamyl alcohol (PCI) extraction. Cells were dissolved in 300μL Tris EDTA (TE) buffer with 300μL PCI solution (phenol pH 6.7 – chloroform – isoamylalcohol 25:24:1) and lysed by micro-bead shearing in a ‘fast prep’ centrifuge (20 sec, 6m/sec)(MP biomedical™). After cell lysis, DNA was isolated and purified using ethanol precipitation. The resulting DNA was diluted to a concentration of 200ng/μL in milliQ H_2_O, based on the DNA concentration measured through absorbance at 260 nm with a NanoDrop spectrophotometer (Isogen).

### Whole genome sequencing and analysis

Genomic libraries were created using the NEBNext^®^ Ultra DNA library Prep Kit for Illumina sequencing (New England Biolabs, US) and genomes were sequenced on an Illumina MiSeq v2 500 (Illumina, US) obtaining a coverage of at least 50x. Standard quality control was performed using FastQC v0.11.7 (69). Paired end reads were aligned using BWA mem v0.7.17 (70) to the annotated genome assemblies of strain B8441 [clade I; Genbank: GCA_002759435.2 (15)] and B11220 [clade II; Genbank: CP043531-CP043537 (26)]. For SNP and Indel identification, the assembly alignment to the annotated genome of strain B8441 (clade I) was used, while CNV analysis was performed using the assembly alignment to reference genome B11220 (clade II) respectively. The genome sequences of all end-point experimentally evolved strains were deposited to NCBI sequencing read archive (SRA) under “BioProject PRJNA664007”. Variants were identified and filtered using GATK v4.1.2.1 (71, 72), with the haploid mode, and including GATK tools HaplotypeCaller and Variant Filtration using “QD < 2.0 ‖ FS > 60.0 ‖ MQ < 40.0”. In addition, variants were filtered if they have minimum genotype (GT) quality < 50, Alternate Allele Frequency < 0.8, or allelic depth (DP) < 10. The final VCF was annotated using SnpEff v4.3T (73). CNVs were identified using CNVnator v0.3 (74), selecting for 1kb genomic windows of significant (*p*<0.01) variation in normalized coverage. The average depth per 5kb window was normalized to the coverage of the whole genome sequence for each isolate and plotted in R (75). Candidate variants were compared with a set of 304 globally distributed *C. auris* isolates representing Clade I, II, III and IV (9).

### PCR and Sanger sequencing

Primers for PCR and Sanger sequencing were designed *in silico* using CLC Genomics Workbench v20.0.3 (digitalinsights.qiagen.com). Primer design was based on B11220 WGS consensus sequences of the regions of interest and sequences of the genes of interest in reference genome *C. auris* B8441 downloaded from the Candida genome database (candidagenome.org). Sequencing primers were designed to include a +/-1000 region of interest (spanning the region with the mutation of interest). All primers are given in **table S1**. Amplification of regions of interest was achieved through PCR using Q5 high-fidelity DNA polymerase (New England Biolabs Inc.). The total reaction volume of 50μL consisted of the 200ng/μL DNA extract, 5μL dNTPs (0.2mM), 10μL Q5 buffer, 0,5μL Q5 polymerase (2 units), milliQ, and 0,4μL of both forward and reverse primer (1 μM). The PCR program consisted of initial denaturation at 98°C for 3 sec, 30 cycles of 98°C for 15 sec, 56°C for 25 sec, 72°C for 2 min and a final elongation step at 72°C for 2 min in a Labcycler Basic thermocycler (Bioké). Correct amplification was verified by performing electrophoresis on a 1% agarose gel at 135V for 25 min. After verification, the sequencing primers (10μM) were added to PCR amplicons and the DNA was sequenced using Sanger sequencing by Eurofins (GATC, Germany).

### Allele-specific PCR (AS-PCR)

The emergence of SNPs and Indels was traced back in whole populations and a maximum of 30 single clones (colonies) per population, using a rapid sequencing free method: allele-specific SNP-PCR. Two primer pairs per gene of interest were designed according to Liu *et al*. (28), consisting of one universal primer or and one mutant-allele primer or wild type-allele primer respectively. Primers consist of an allele specific region at the 3’ terminal nucleotide of the mutant or wild type allele specific primer. Additionally, a mismatch at the 3th nucleotide from the 3’ terminal was included to increase annealing specificity (28). All primers used for AS-PCR are listed in **table S1**.

To validate primer specificity, a temperature gradient PCR was performed in which annealing temperature varied between 60°C and 70°C. AS-PCR sensitivity was assessed by performing PCR on serial dilution of reference DNA template. All PCR reactions were performed in a total reaction volume of 20μL consisted of 1μL of 1/20 dilution of the pure PCI DNA extract, 5μL dNTPs (0.2mM), 10μL TaqE buffer, 0,5μL TaqE polymerase (2 units), milliQ, and 0,4μL of both forward and reverse primer (1 μM). The PCR program consisted of initial denaturation at 98°C for 3 sec, 30 cycles of 98°C for 15 sec, 56°C for 25 sec, 72°C for 2 min and a final elongation step at 72°C for 2 min in a Labcycler Basic thermocycler (Bioké). Amplification and thus the presence or absence of a mutation was verified by performing electrophoresis on a 1% agarose gel at 135 V for 25 min.

### Gene expression and copy number variation analysis

*C. auris* cells from a single colony grown overnight on YPD agar (2% glucose) were enriched in RPMI-MOPS (2% glucose) medium for 16h. These cultures were diluted to 10^8^ cells in a volume of 50ml fresh RPMI-MOPS (2% glucose) medium and incubated for 8h at 37°C in a shaking incubator to ensure the harvested cells are growing in the exponential growth phase. Next, cells were harvested by centrifugation, washing in ice cold PBS, snap freezing in liquid nitrogen and stored at −80°C.

For gene expression analysis (RNA extraction and RT-qPCR), cells were resuspended in 1ml trizol and lysed by micro-bead shearing in a ‘fast prep’ centrifuge (20 sec, 6m/sec)(MP biomedical™). Nucleotides were extracted by washing the lysate supernatant with chloroform (360μL) and isopropanol (350μL) and precipitated by washing three times with 70% ethanol. Nucleotide concentrations and purity were measured spectrophotometrically using a NanoDrop ND-1000 (Isogen Life Science). Extracts were diluted to 1μg pure nucleotide concentration and purified by DNase treatment (New England Biolabs). cDNA was synthesized from RNA by using the iScript cDNA synthesis kit (Bio-Rad) according to the manufacturer’s recommendations. Real Time qPCR was performed using GoTaq polymerase (Promega) and the StepOnePlus real-time PCR thermocycler (ThermoFisher) as follows: activation at 95°C for 2 min, 40 cycles of denaturation at 95°C for 3 seconds and annealing/extension at 60°C for 30 seconds. Primers used for qPCR were designed with the PrimerQuest tool of IDT (https://eu.idtdna.com/Primerquest/) and are listed in **table S1**. A total of 8 housekeeping genes involved in various cellular processes were assessed, of which the 3 most stable candidates were used in the analysis (*ACT1, LSC2, UBC4*). Gene expression analysis was performed using qBasePlus software. Fold Change (with SD) was plotted from log2(Y) transformed data and compared statistically (using a one-way ANOVA with multiple comparisons in respect to wt) with GraphPad Prism. Expression analysis in each strain was performed using 3 biological repeats each represented by the average of 2 technical repeats.

For copy number variation analysis, gDNA was extracted as described in ‘DNA extraction’ of methods section and standardized concentrations of 0.5 ng/μL of gDNA were used to quantify target markers (*TAC1b* and *ERG11* respectively) by qPCR using the same primers, protocol and analysis as described above.

## Acknowledgement

This work was supported by an FWO personal research grant nr. 11D7620N awarded to H.C. A travel grant awarded by FWO allowed a fruitful research stay of H.C. at Broad Institute (Boston, US). C.A.C and J.F.M. were supported by the National Institute of Allergy and Infectious Diseases, National Institutes of Health, Department of Health and Human Services, under award U19AI110818 to the Broad Institute. We want to thank the group of prof. Steenackers (KULeuven) for their advice on experimental evolution.

---

**Figure S1.**
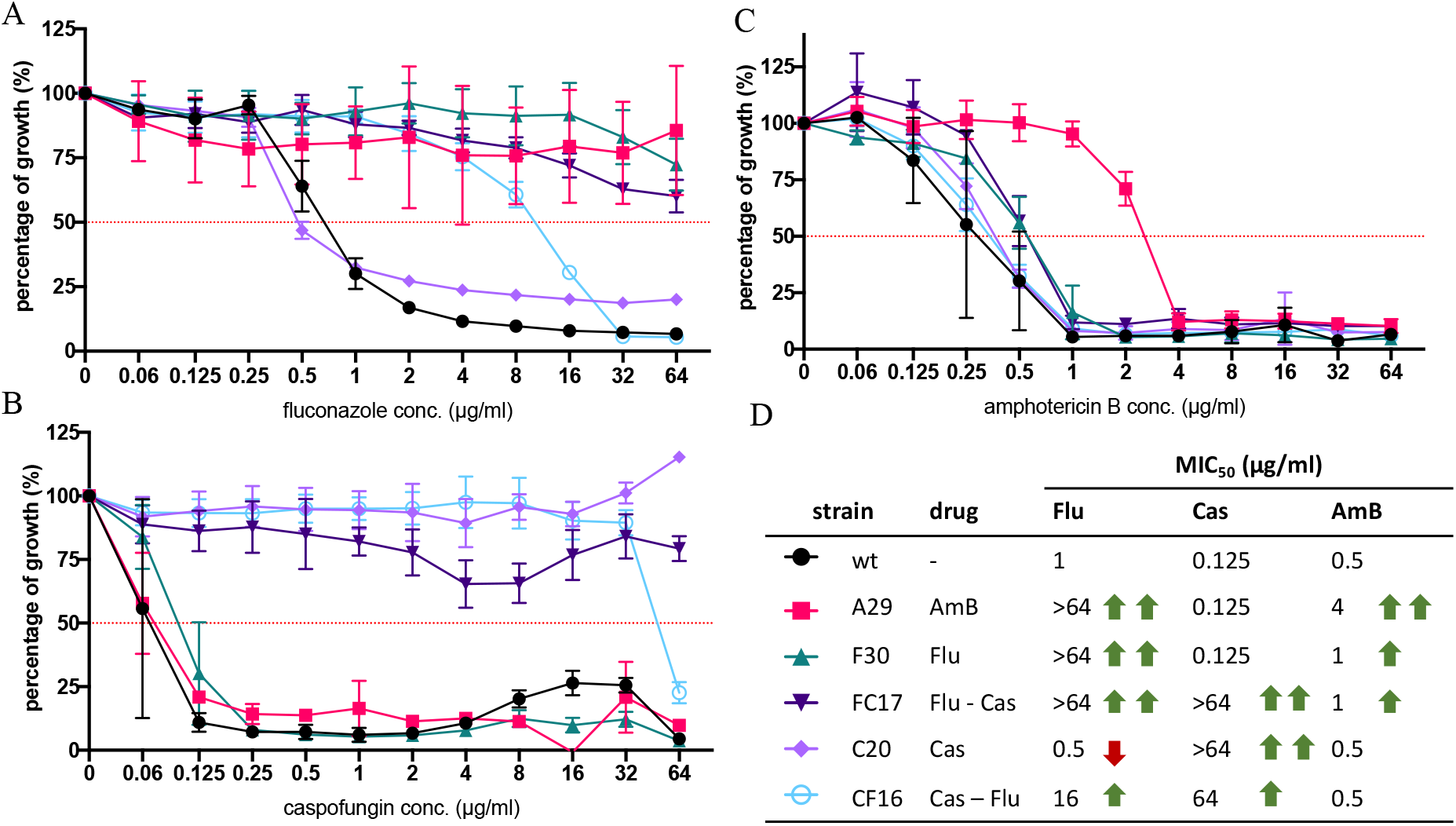
Resistance profiles of end-point evolved strains. **A-C)** Growth profiles in a BDA of the end-point evolved strains (A29, F30, FC17, C20, CF16) and control strain (wt) for fluconazole **(A)** caspofungin **(B)** and amphotericin B **(C)** respectively. Percentage of growth was calculated from growth without drug and based on OD_600_ measurements after 48h of incubation at 37°C. Each data point and its standard deviation is calculated from 3 biological repeats, each represented by the mean of 2 technical repeats. **D)** Summary of MIC_50_ values for each strain and indication of (relatively low 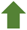/high 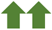) increase or decrease 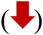 in MIC in respect to parental strain (wt) MIC.

**Figure S2.**
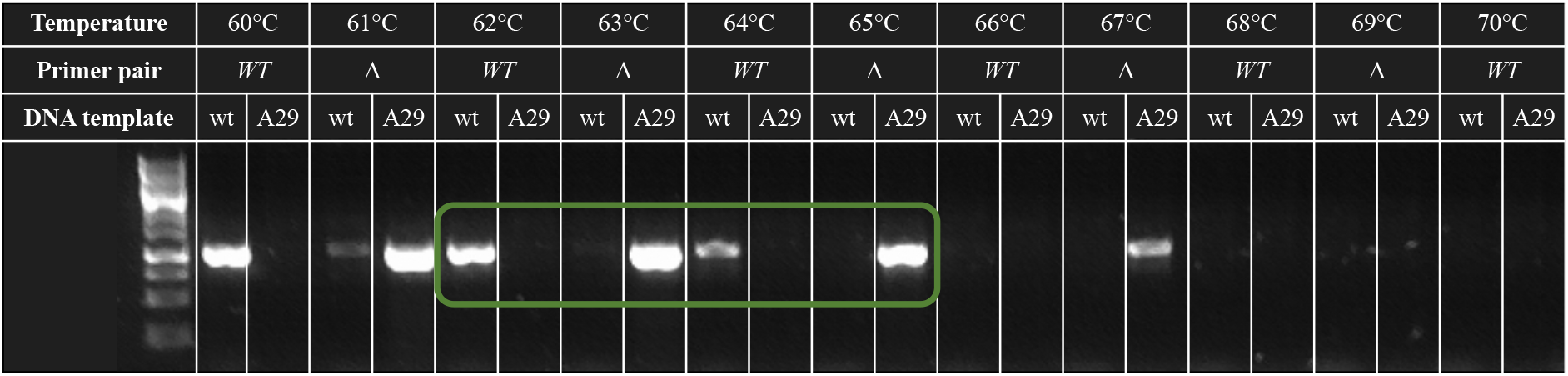
An example of the temperature specificity range of allele-specific primers. Here a temperature gradient PCR was performed using the *MEC3* allele-specific primers (shown in supplementary table S1) on gDNA of the wild type strain (wt) and strain A29, containing the gCg/gTg|A272V mutation in *MEC3* (see Table 1). Primer pairs ‘*WT*’ and ‘Δ’ indicate the primers targeting the wild type allele (*CauMEC3_B11220_PCR/Seq1_F* and *CauMEC3_SNP_A272A_R*) and mutated allele (*CauMEC3_B11220_PCR/Seq1_F* and *CauMEC3_SNP_A272V_R*) respectively and are given in supplementary table S1. The green grid indicates a primer specific temperature range (63-64°C resp.)

**Figure S3.**
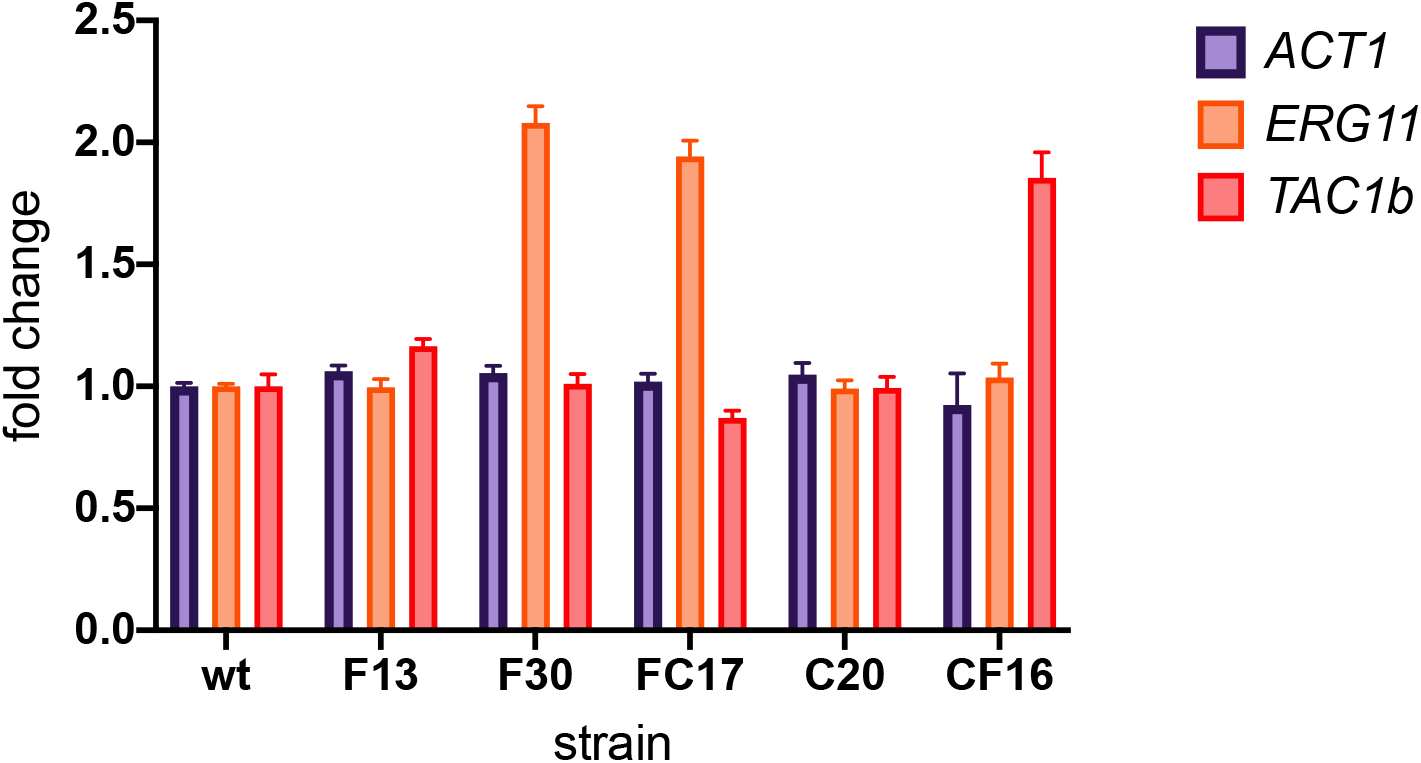
**Copy number variation quantification** of *TAC1b* (marker on the Chr5 duplication in CF-lineage, see figure 6), *ERG11* (marker on the segmental duplication of Chr1 in F- and FC-lineage, see figure 6) and *ACT1* (reference) by qPCR on gDNA. CNVs were determined for 1 biological repeat represented by 3 technical repeats.

**Table S1.**
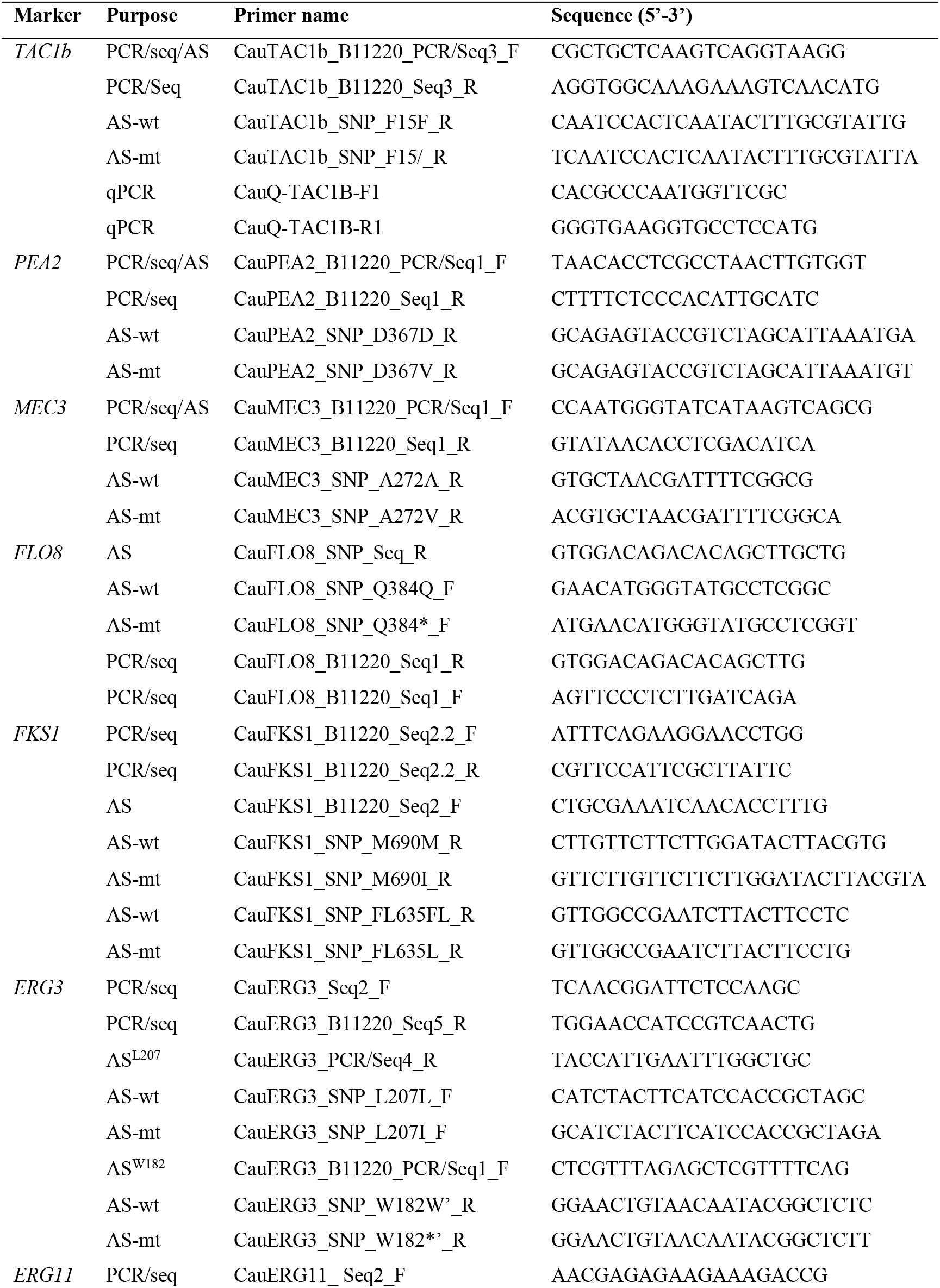
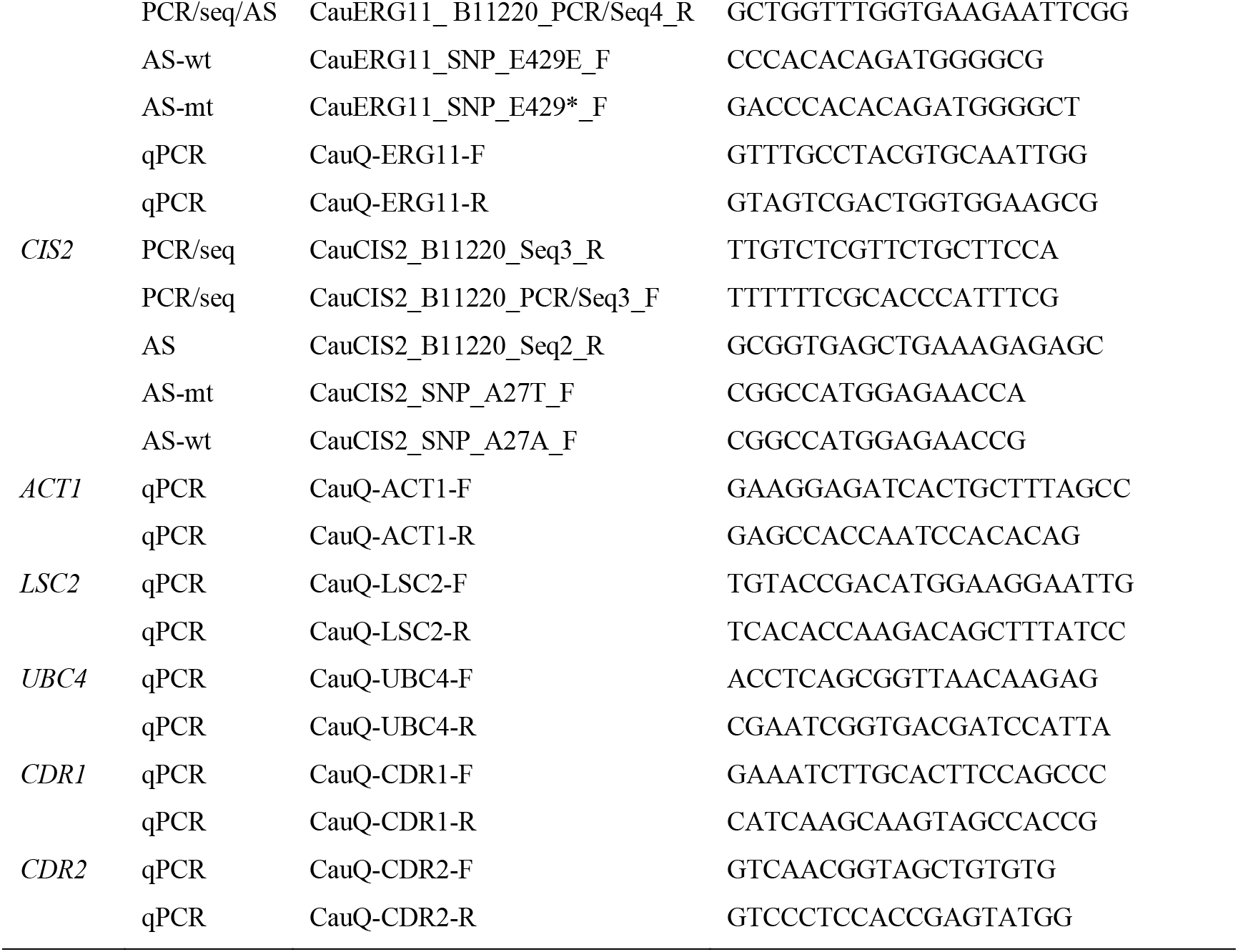
All primers used in this study. Primers are arranged per marker. ‘Purpose’ indicates whether the primer was used for PCR and sequencing (PCR/seq), CNV or expression analysis (qPCR) or allele-specific PCR (AS). Primer pairs for AS-PCR consist of a universal PCR-sequencing primer (indicated by ‘PCR/seq/AS’) or universal AS-primer (indicated by ‘AS’) and one allele-specific primer (indicated by ‘AS-wt’ for the wild type allele and ‘AS-mt’ for the mutant allele).

